# The molecular basis of immunosuppression by soluble CD52 is defined by interactions of N-linked and O-linked glycans with HMGB1 Box B

**DOI:** 10.1101/2024.10.24.620132

**Authors:** Nicholas J. DeBono, Silvia D’Andrea, Esther Bandala-Sanchez, Ethan Goddard-Borger, Muhammad A. Zenaidee, Edward S.X. Moh, Elisa Fadda, Leonard C. Harrison, Nicolle H. Packer

## Abstract

Human soluble CD52 is a short glycopeptide comprising 12 amino acids (GQNDTSQTSSPS) which functions as an immune regulator by sequestering the pro-inflammatory high mobility group box protein 1 (HMGB1) and supressing immune responses. Recombinant CD52 has been shown to act as a broad anti-inflammatory agent, dampening both adaptive and innate immune responses. This short glycopeptide is heavily glycosylated, with a complex sialylated N-linked glycan at N3 and reported O-linked glycosylation possible on several serine and threonine residues. Previously we demonstrated that specific glycosylation features of CD52 are essential for its immunosuppressive function, with terminal α-2,3-linked sialic acids required for binding to the inhibitory SIGLEC-10 receptor leading to T-cell suppression. Using high resolution mass spectrometry, we have further characterised the N- and O-linked glycosylation of Expi293 recombinantly produced CD52 at a glycopeptide and released glycan level, accurately determining glycan heterogeneity of both N- and O-linked glycosylation, and localising the site of O-glycosylation to T8 with high confidence and direct spectral evidence. This detailed knowledge of CD52 glycosylation informed the construction of a model system, which we analysed by molecular dynamics simulations to understand the mechanism of recognition and define interactions between bioactive CD52, HMGB1 and the SIGLEC-10 receptor. Our results confirm the essential role of glycosylation, more specifically hyper-sialylation, in the function of CD52, and identify at the atomistic level specific interactions between CD52 glycans and the Box B domain of HMGB1 that determine recognition, and the stability of the CD52/HMGB1 complex. These insights will inform the development of synthetic CD52 as an immunotherapeutic agent.

## Introduction

Human CD52 is a GPI-anchored glycopeptide composed of 12 amino acids (GQNDTSQTSSPS) in its released soluble form (1). Despite its small size, this peptide has previously been shown to carry both N- and O- linked glycosylation (2, 3). Our previous studies (4) identified soluble CD52 as having broad immunosuppressive properties. Activated T cells release CD52 which, after sequestering HMGB1, binds to the inhibitory sialic acid-binding immunoglobulin (Ig-like) lectin-10 (SIGLEC-10) on T cells (2, 5). HMGB1 is comprised of two structurally identical domains, Box A and Box B; CD52 interacts with Box B2, which when uncomplexed is pro-inflammatory. The immunosuppressive properties of CD52 identify it as a potential immunotherapeutic agent.

Previous work on the characterisation of CD52 has identified two glycans on the peptide, an N-linked glycan at N3, and an O-linked glycan at several possible O-glycosites, at T8, S9 and/or S10 (3). These glycosylations are highly complex in terms of both their heterogeneity and overall glycan structure, and act as the main feature for interaction between CD52, HMGB1-Box B and SIGLEC-10 (3). The structure of CD52 glycans has remained an ongoing analytical challenge, with its complexity and site of O-glycosylation being only partially solved (1, 3, 6). Glycans on CD52 contain a high abundance of α-2,3 sialyl glycoforms, which we confirmed are essential for its immunosuppressive activity (3).

However, the relevance of sialylation has not been confirmed for any O-glycosylation present on bioactive CD52. Previously optimised methods for production of recombinant soluble CD52 ^4^ produce very low yields of the active glycopeptide (3). As glycosylation is required for activity of CD52, the glycoforms of active CD52 need to be fully characterised for development of CD52 as an immunotherapeutic agent. Here, we extend previous work characterising the recombinant CD52 glycome by defining both the N- and O-linked active rCD52 glycoforms in greater detail, including N- and O-glycan structural characterisation, and by determining both the site and occupancy of O-glycosylation. These experimental results then informed modelling to explain at the atomistic level how glycans mediate the interactions between CD52 and HMGB1-Box B, leading to recognition by SIGLEC-10. More specifically, we found through extensive model sampling that hyper-sialylation of both O- and N-glycans on CD52 determines its conformational propensity and allows the glycopeptide to engage with specific contact amino acid residues in HMGB1-Box B. These results explain the preference of CD52 for the HMGB1-Box B domain over the structurally identical Box A. Ultimately, the structure of the CD52/HMGB1-Box B complex we identified through Molecular Dynamics (MD) simulations favours the exposure of the CD52 N-glycans sialylated arms towards the solvent, in a conformation accessible for recognition by SIGLEC-10.

## Results

### Recombinant CD52 displays glycan heterogeneity and a low abundance of bioactive forms

When soluble CD52-Fc fusion protein was fractionated by anion exchange chromatography, only a minority of individual fractions exhibited immunosuppressive activity (3). Activity increased with increasing negative charge, before decreasing rapidly (Figure 1A). The most active fraction, #34, comprised only 0.75 % of the total CD52-Fc protein produced.

**Figure 1:**
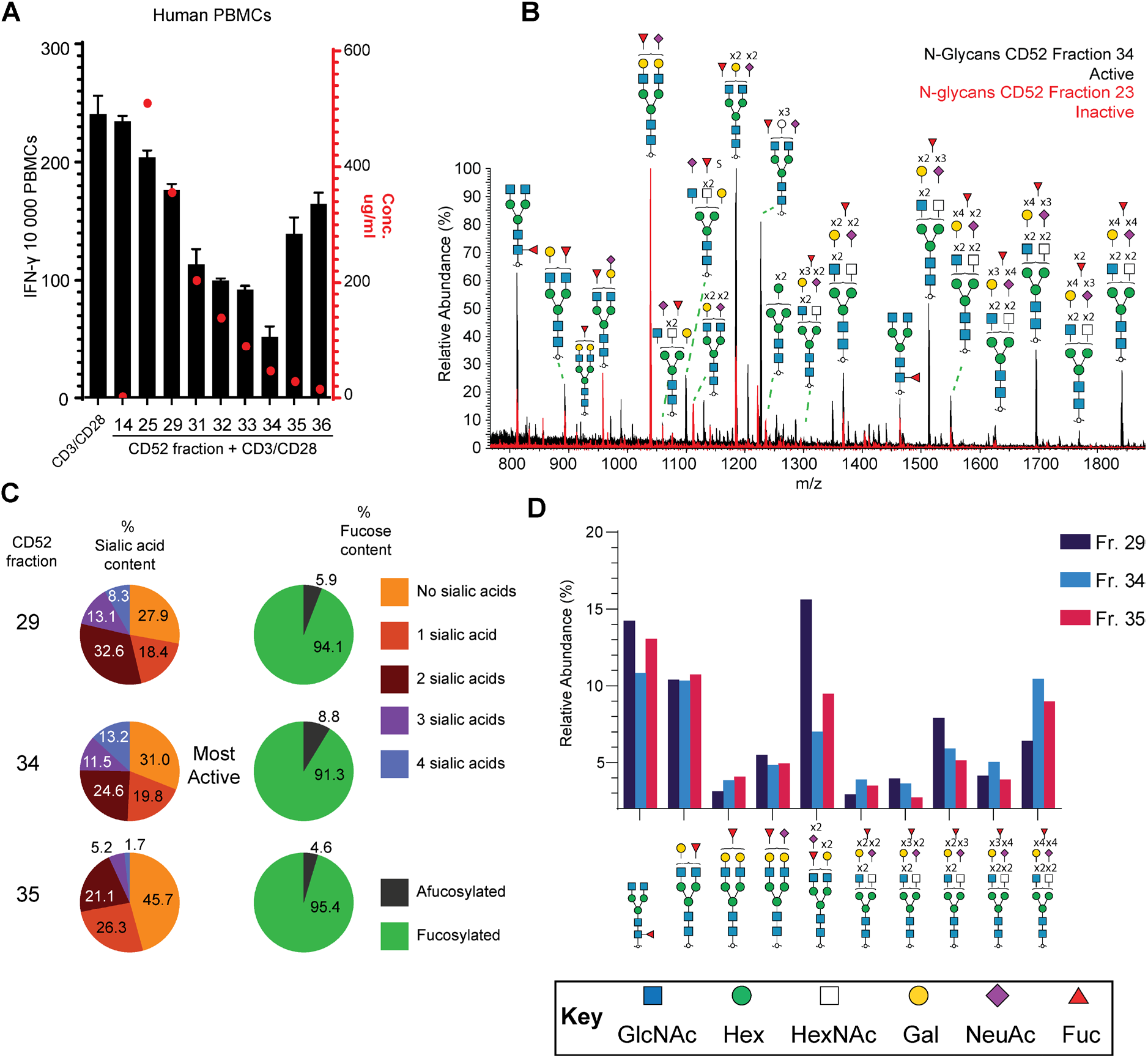
Characterization of the immunosuppressive activity and N-glycan compositions of recombinant CD52-Fc fractions. A) Bars: IFN–γ ELISpot assay of CD52-Fc activity after fractionation by anion exchange chromatography (representative fractions shown). 10,000 peripheral blood mononuclear cells (PBMCs) were incubated in 200 μL per well either with or without anti-CD3/CD28 Dynabeads (1 bead/cell) in the presence of CD52-Fc fractions (F14–F36) at 5 ng/ml. The left Y-axis shows the number of IFN-γ-positive ELISpots, a decrease indicating immunosuppressive activity of CD52-Fc. The red dots correspond to the protein concentration (µg/ml) of CD52-Fc in each fraction, measured by Qubit absorbance, with the right Y-axis referencing this data. B) Summed MS1 spectrum of the active fraction 34 (black) is overlaid with the spectrum of the most abundant fraction by protein weight, #23 (red), which is inactive. A greater proportion of highly sialylated N-linked compositions can be seen in the active fraction 34. C) % Sialic acid (left) and % fucosylation (right) content of N-glycans in selected anion exchange CD52-Fc fractions. The highly active fraction 34 contains a high abundance of highly sialylated N-glycans, while all fractions are heavily fucosylated. D) The top 10 N-glycan compositions with more than 2.5% relative abundance in active fraction 34, compared to two other anion exchange fractions and organised by increasing *m/z* (M-H). All compositions have been shown according to SNFG cartoon nomenclature guidelines(7).

### N-glycosylation on Asparagine 3 of CD52-Fc is highly heterogeneous and highly sialylated

After removal of the Fc tag on CD52 by cleavage with Factor Xa (FXa), N-glycans were released from anion exchange fractions of CD52, cleaned and analysed via negative mode PGC ESI-LC tandem mass spectrometry. 192 unique glycan structures were completely or partially determined from 80 glycan compositions using previously reported diagnostic MS2 ions and reported retention time rules of PGC separation (Supporting information S1) (8–13). Almost all N-glycans characterised on CD52-Fc were fucosylated, predominantly with a core fucose motif (Figure 1C and D, Supporting information S1). When examining the comparative N-glycome of all fractions, an increase in degree of sialylation could be observed as fractions increased their elution from anion exchange (Figure 1B), with 24.7 % of the N-glycan population of the most active fraction 34 containing at least three sialic acids (Figure 1C, Supporting information S1). Interestingly, the N-glycome of the relatively inactive fraction 35, collected directly after the most active fraction 34 (Figure 1A), only carried a 6.9 % abundance of N-glycans with at least three sialic acids and contained 45.7 % unsialylated N-glycans (Figure 1C, Supporting information S1). We have previously shown that sialylation, particularly 2,3 linked, is essential for the activity of CD52-Fc (3). Of the 10 most abundant N-glycan compositions observed on the active rCD52 fraction, seven were sialylated, usually with two or more sialic acids (Figure 1D).

When comparing fractions, the tetra-sialylated composition Hex7HexNAc6Sial4Fuc1 was present in highest relative abundance in the active fraction 34 (Figure 1D, Supporting information S1).

Previously, we observed polyLacNAcs on native soluble CD52 N-glycans from spleen (3). Analysis of the tandem mass spectra of released N-glycans from CD52-Fc also showed some polyLacNAc N-glycan extensions (Supporting Information S2).

Following FXa cleavage and N-glycan release, O-glycan alditols were chemically removed from CD52 fractions using reductive β-elimination and analysed using well established PGC-ESI tandem mass spectrometry (8, 14, 15). Eight O-glycan structures were observed from six compositions, with the (Hex2HexNAc2NeuAc2, GlyToucan ID G52120NK) occupying 55.5 % of the O-glycome of the most active fraction 34. Both observed doubly sialylated abundant O-glycans were only present in one isomeric form indicating one type of sialic acid linkage, most likely α2,3 (Figure 2B, Supporting Information S3). Double sialylation of the O-glycans on CD52 appeared to be conserved across less active, later eluting fractions (Fractions 23 to 35), with the second most abundant O-glycan as the doubly sialylated mucin-type core 1 structure (HexHexNAcNeuAc2, GlyToucan ID G63110FE) present at an average 17.6 % relative abundance (Figure 2A, SupportingSupporting Information S3).

**Figure 2:**
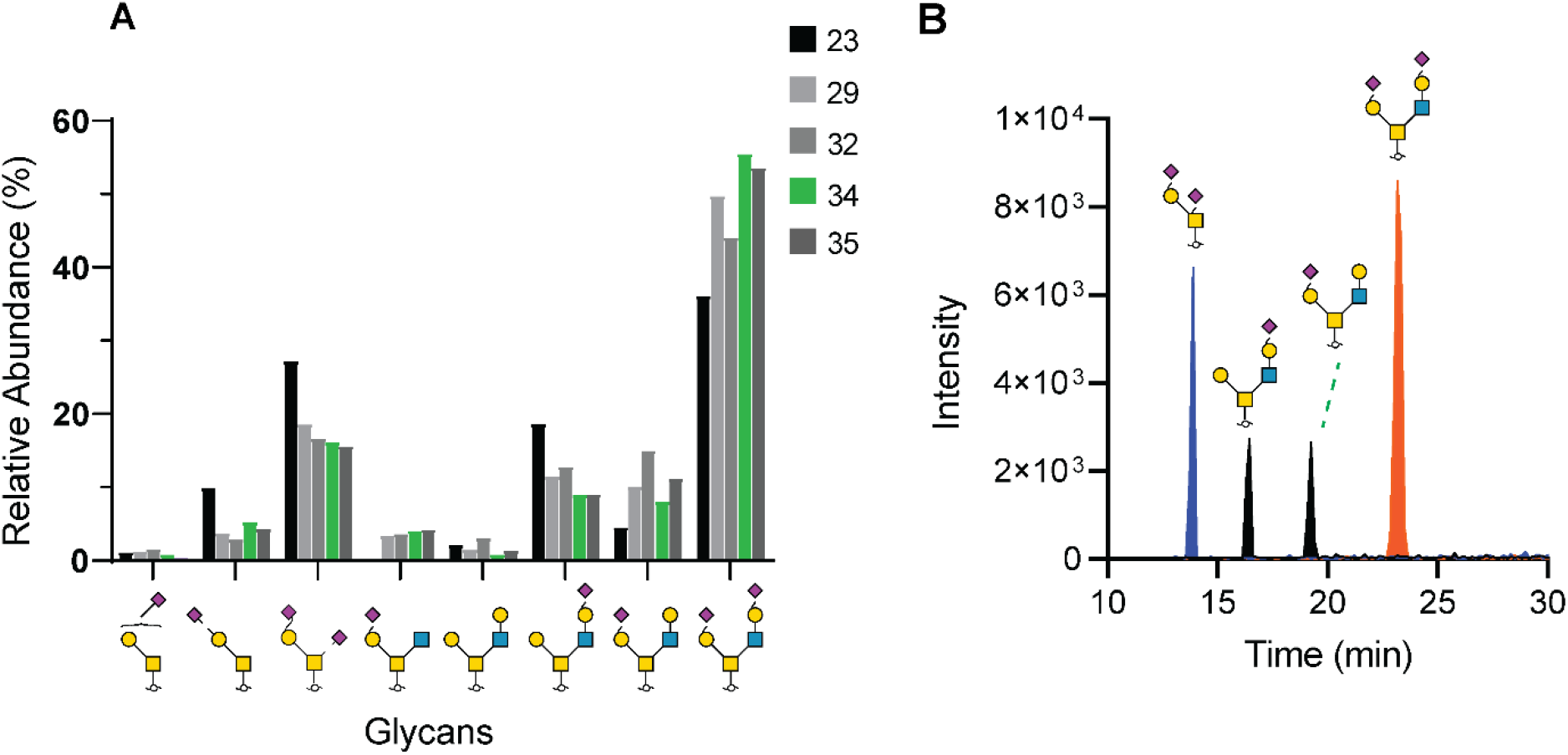
Characterization of O-glycan structures and sialic acid linkage variants in active CD52 fraction 34. A) O-glycan structures identified in specific CD52 fractions, with the most active fraction 34 highlighted in green. Fraction 34 exhibited the highest relative abundance of di-sialylated core 2 O-glycan (GlyToucan ID G52120NK). B) Extracted ion chromatograms displaying the three most intense O-glycan mass compositions in fraction 34. Both doubly sialylated O-glycans appeared as a single isomer, suggesting only one type of sialic acid linkage, while glycans containing one sialic acid appeared as two unique isomers, due to alternative arm sialic acid linkage addition.

### O-glycosylation of rCD52 is localised to Threonine 8 with low site occupancy

The site of O-glycosylation was then determined on CD52, after treatment with FXa to remove the Fc fusion and treatment with PNGase F to remove N-glycans, using a glycopeptide diagnostic ion triggered EtHcD LC-MSMS workflow. Treatment with FXa of the CD52-Fc results in a 29-mer glycopeptide (Supporting Information S10), which was used in subsequent searches. All fractions were analysed using the glyco-O-Pair default settings within Fragpipe in MSFragger software (16, 17), with the identified six compositions of the O-glycome as the glycan search space. As previous attempts to confidently identify the O-glycosite of CD52 have resulted in ambiguity (3), we chose a very strict cut off of reported confidence level (level one only) and an O-Pair score (>20) to ensure accurate determination of the O-glycosite. Using these filters within O-Pair, 47 glycoPSMs were identified (Supporting Information S5), which were then manually confirmed for the presence of peptide ions fragments on either side of the predicted site of glycosylation. All filtered glycoPSMs localised the site of O-glycosylation to T8 (Figure 3), with the most highly confident spectral matches occurring with non- or partially-sialylated O-glycopeptides (Supporting Information S5). Comparing the proportion of O-glycosylated to non-O-glycosylated CD52, site occupancy at the T8 O-glycosite was calculated to be just 7.5 % of analysed CD52 peptides across all fractions (Supporting Information S6). This O-glycosite determination aligns with glycosite predictions by the O-glycosylation predictive software IsoGlyP (18), which predicts T8 as the most likely site for O-glycosylation by all modelled transferases (Supporting Information S7). By contrast, the N3 N-glycosite of CD52 was found to be occupied, on average, 99% of the time across all fractions on CD52 when comparing non-N-glycosylated CD52 to the deamidated CD52 peptides pre- and post-N-glycan removal by PNGase F digestion (Supporting Information S6).

**Figure 3:**
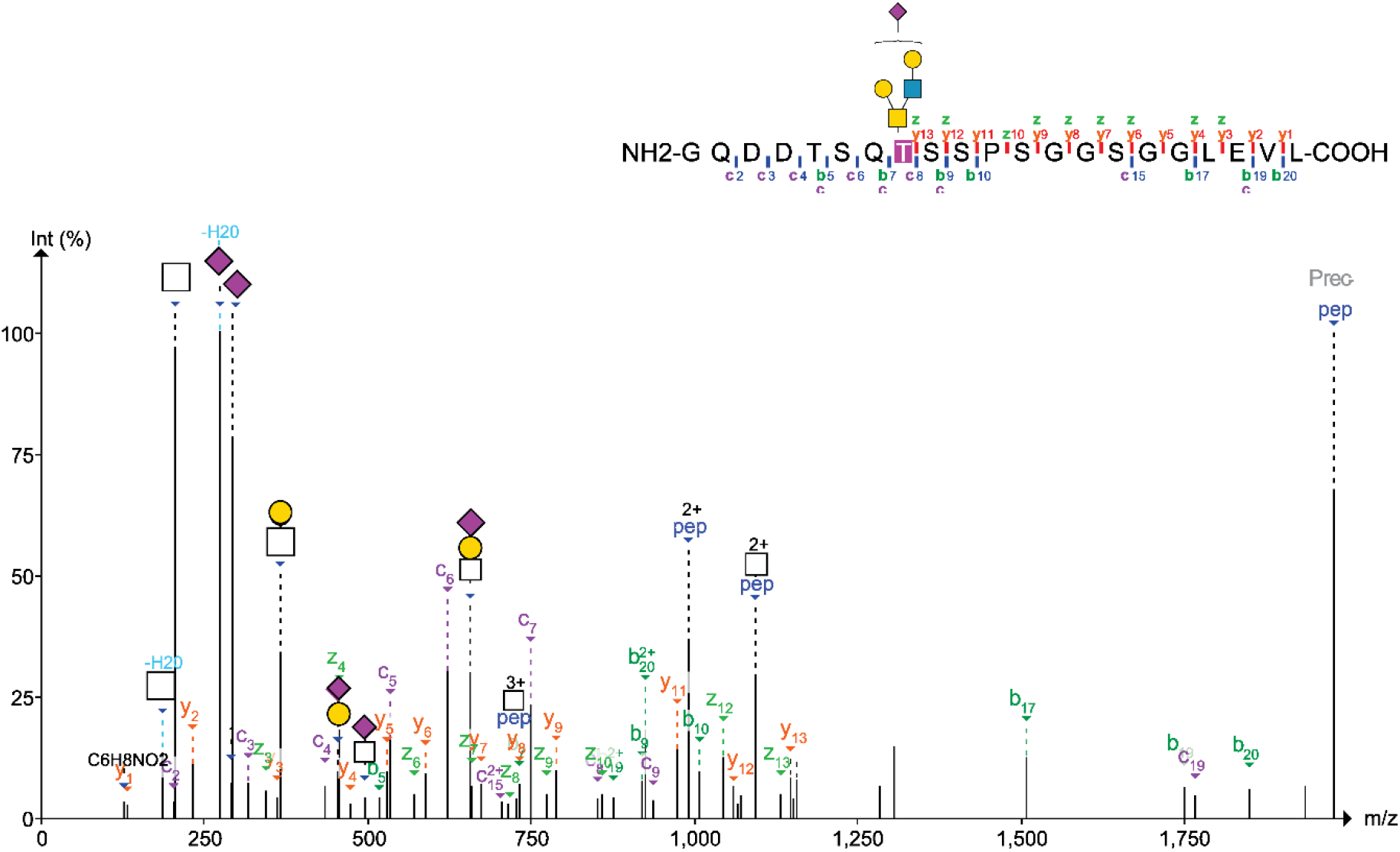
Example of annotated spectra for O-glycosite determination, showing a paired EtHcD scan that includes both b/y and c/z peptide fragments. Glycan diagnostic oxonium ions are shown with glycan fragment mass.

### Molecular dynamics simulation reveals an intricate interaction between glycosylated CD52 and HMGB1 Box B

We used an approach based on deterministic sampling by molecular dynamics (MD) simulations to understand the role of sialylation on the recognition and binding of the 12 residue CD52 glycopeptide (aa 25-36 region from the Uniprot P31358 CAMPATH-1 antigen with sequence GQNDTSQTSSPS) to the HMGB1 Box B. We first analysed the effects of sialylation on the conformational equilibrium of the isolated CD52 glycopeptide by running two sets of five uncorrelated MD simulations, in one set the peptide was linked to the fully sialylated N- and O-glycans (GlyToucan ID’s G80552MJ and G42089IU respectively) based on the analytical glycoanalysis described above and in the other, the peptide was linked to the same glycan structures but without terminal sialic acids (GlyToucan ID’s G56655CC and G96017QA respectively) (Figure 4a and 4b). The analysis of the CD52 structures obtained from 10 μs of cumulative sampling, 5 μs for each set, showed that hyper-sialylation stabilises an extended structure of the peptide, by means of conformational changes in the torsion angles values of the backbone at, or in the immediate vicinity of, the glycosylation sites. More specifically, the Ramachandran analysis of the conformation of the CD52 glycopeptide shows significant changes in the conformational propensity at bonds 2 and 8, corresponding to the peptide bonds centred at N3 and S9, respectively (Figures 4c and 4d). In the case of N3 (bond 2), the conformational preference changes from an extended β strand conformation in the presence of a core-fucosylated tetra-antennary and fully sialylated *N-*glycan (GlyTouCan ID G80552MJ) to a 3-10 helical turn when the same *N-*glycan structure does not have terminal sialic acids (GlyTouCan ID G56655CC). The absence of sialylation on the extended core 2 *O-* glycan (GlyTouCan ID G96017QA) at T8 stabilises the localised propensity for a left-handed α helical conformation, which extends to S9. The increased helicity of the CD52 glycopeptide in the absence of sialylation ultimately leads to a more compact structure, as shown also by the analysis of the radii of gyration (Rg) (Figure S7). The Ramachandran plots for all the peptide bonds in the presence and absence of sialylation are shown in Figures S9 and S10, respectively.

**Figure 4.**
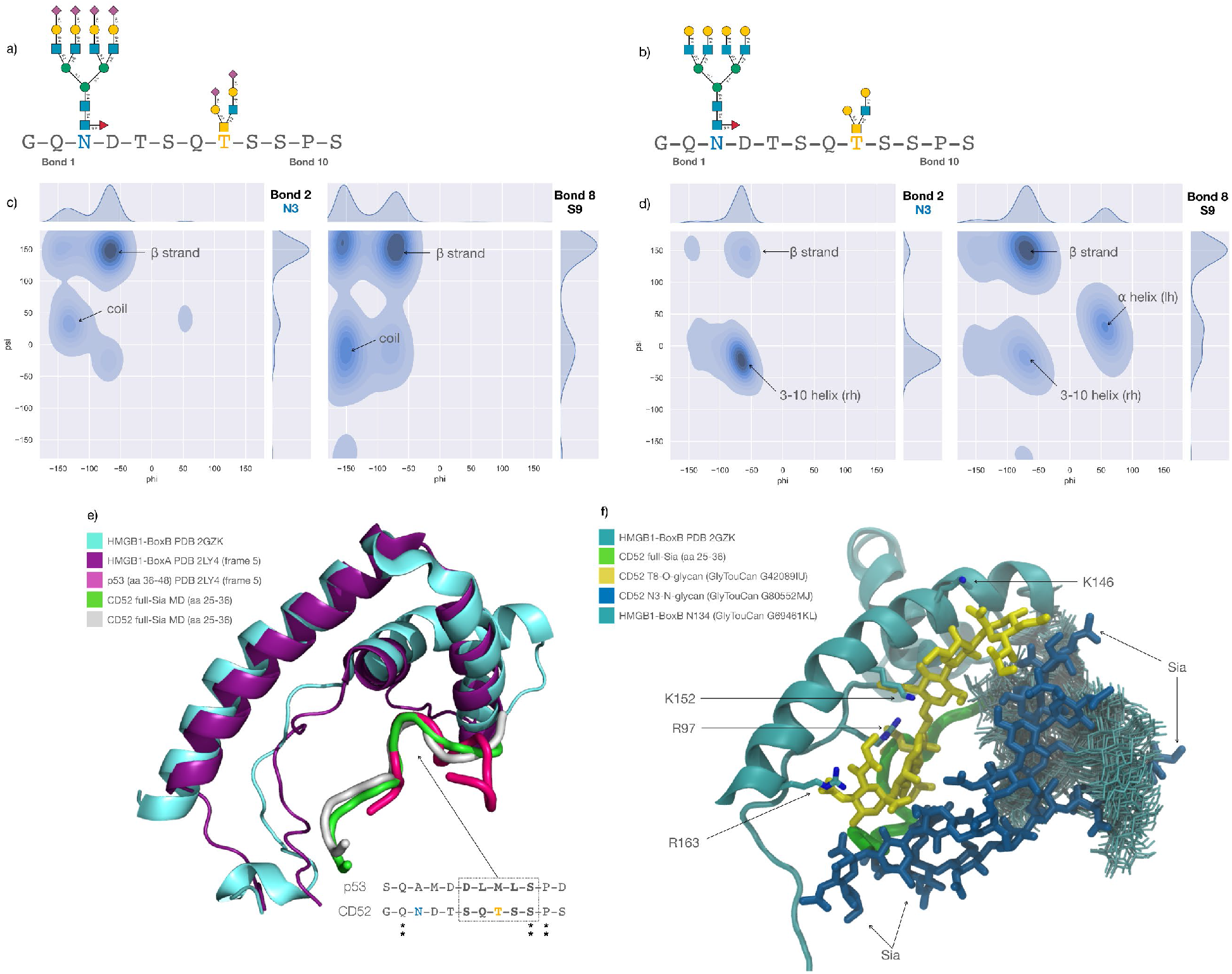
Structural and dynamic analysis of CD52 glycopeptide variants and their interaction with HMGB1 domains. A) Sequence of the 12 residue CD52 active peptide (aa 25-36) with the hyper-sialylated glycan structures at N3 (GlyTouCan ID G80552MJ) and T8 (GlyToucan G42089IU) depicted using SNFG nomenclature. B) Sequence of the 12 residue soluble CD52 peptide (aa 25-36) with the non-sialylated glycan structures (GlyTouCan ID G56655CC and G96017QA) at N3 and T8, respectively. C) Ramachandran plot of peptide bonds 2 and 8, centred at N3 and S9, respectively, obtained from 5 μs of cumulative MD sampling of the hyper-sialylated CD52 glycopeptide. The secondary structure types associated with torsion angles are indicated. Kernel Density Estimates (KDE) are shown on the top and right of the plot. D) Ramachandran plot of the peptide bonds 2 and 8, centred at N3 and S9, respectively, obtained from the 5 μs of cumulative sampling of the non-sialylated CD52 glycopeptide, with corresponding KDE values on above and on the right-hand side of the plot. All plots were made with seaborn (https://seaborn.pydata.org/). E) Structural alignment of the HMGB1-Box A (purple cartoon representation, PDB 2LY4) bound to p53 (pink cartoon representation, PDB 2LY4) to the HMGB1-Box B (cyan cartoon representation, PDB 2GZK) with two selected representative snapshots (green and white cartoon representation) from the cumulative 5 μs MD simulation of the hyper-sialylated CD52 glycopeptide. Sequence alignment of the HMGB1- Box A-bound p53 peptide to the active CD52 glycopeptide where the sequence corresponding to the helical turn is highlighted in the dotted box. The T8 of CD52 is highlighted in yellow. Graphical rendering and sequence alignment with pymol (https://pymol.org/2/). F) Snapshot (frame 200 ns) from the 1 μs MD trajectory of the reconstructed 3D model of HMGB1-BoxB (cyan) in complex with the hyper-sialylated CD52 glycopeptide (green). The N- and O-glycans are shown with sticks in blue and yellow, respectively. The key protein residues identified to engage in interactions with the O-glycan at T8 are indicated with labels and shown with sticks. The position of the terminal sialic acids of the CD52 N-glycan are also highlighted with labels. The conformation of the HMGB1-Box B N-glycan is shown with multiple snapshots collected every 20 frames (1 ns each) through the duration of the trajectory. Graphical rendering with VMD (https://www.ks.uiuc.edu/Research/vmd/). All cartoon representations of glycans shown according to SNFG symbol nomenclature guidelines, with key in figure 1.

To our knowledge, at this time there is no available structural information on the complex between CD52 and HMGB1 Box B, or on the basis for the preference of the CD52 glycopeptide for Box B relative to the structurally identical domain ^2^, Box A, to which CD52 does not bind. In order to gain insight on the molecular basis for the preferential recognition of hyper-sialylated CD52 glycopeptide for HMGB1 Box B, we analysed the structure of the complex between the HMGB1 Box A and a section of the p53 transactivation domain (19) (aa 36-48). Although the functions of these two complexes are entirely unrelated, the structural features of the HMGB1 Box A/p53 complex provides important clues that ultimately led us to build a working model for our complex. The rationale behind choosing the HMGB1-BoxA/p53 as a template is based on two main considerations. First, HMGB1 Box A and Box B share a high structural similarity, with a backbone RMSD value of 2.27 Å over 54 atoms, and a sequence identity of 25.33% for the human HMGB1 (Uniprot P09429) (20) (Figure 4e). Second, the structural alignment of representative snapshots from the MD simulations of the CD52 hyper-sialylated glycopeptide to the Box A-bound p53 conformation indicates a keen similarity between the two peptides around the O-glycosylation site on CD52 at T8 (Figure 4e). Guided by these considerations, we generated a 3D model of the HMGB1-BoxB in complex with the CD52 glycopeptide, in which glycosylation on the peptide and on the HMGB1-BoxB at N134 was restored with GlycoShape (21) to complement the tight steric requirements.

The most striking and unique feature of this 3D model is that the main contacts occur between the HMGB-1 and the sialylated core 2 O-glycan, which directly faces the Box B surface, and not between HMGB1 Box B and the CD52 peptide (Figure 4f). Also, in this model the large tetra-antennary N-glycan at N3 is orientated with the arms directed towards the solvent, and thus accessible for binding (potentially to SIGLEC10 (22)). We assessed the stability of this complex through a single trajectory of MD simulation with a 1 μs production phase ran all degrees of freedom unrestrained. The simulation allowed us to identify a set of key residues of HMGB1-BoxB that make specific contacts with the fully sialylated core 2 at T8, namely R97, K146, K152, R163, see Figure 4f. Interestingly, all of these residues, except K152, are not conserved in HMGB1-BoxA, where R97 is G, K146 is D and R163 is I, providing a rationale for the preference of the hyper-sialylated peptide for Box B and not Box A

Ultimately, aside from other considerations and based on this 3D model of the complex alone, lack of sialylation of the core 2 O-glycan at T8 would prevent binding of CD52 to HMGB1 Box B as most contacts with the protein involve directly the terminal sialic acids (Figure 4f). The MD simulation also shows that the terminal sialic acids on the CD52 *N-*glycan are mostly orientated towards the solvent and that the N-glycan on HMGB1-BoxB wraps around the CD53 N3 N-glycan. As a note of caution, additive force fields tend to enhance glycan-glycan interactions (23), so the extent of these contacts may be overestimated.

## Discussion

In this study, we have extended previous work to determine the structure of active immunosuppressive CD52, produced recombinantly (CD52-Fc) (3), further characterising both the complex, heavily sialylated N-and O-glycome. Glycosylation is a critical requirement for both CD52 activity (3) and SIGLEC-10 interaction (22), in both cases 2,3-sialylation conferring greater bioactivity than 2,6 sialylation. N-glycans identified in active fractions of CD52-Fc were overwhelmingly core fucosylated and heavily sialylated, with N-glycans containing at least two sialic acids comprising 49.3 % of the relative abundance of all observed N-glycans on active CD52-Fc. PolyLacNAcs were also present, as in native soluble spleen CD52 (3). Elongated linear and branched polyLacNAc chains are ubiquitiously expressed in normal human adults (24), and have been broadly implicated in biological processes (25–29); however, their direct influence on glycan binding is relatively poorly understood due to inability to confidently resolve individual polyLacNAc isomers (25). In the context of soluble CD52, we propose that the presence of linear polyLacNAc structures ending in a terminal sialic acid could effectively enables deeper interaction between CD52 N-glycan sialic acids and the immunoinhibitory SIGLEC-10.

Compared to the N-glycome, the O-glycome observed on CD52 had low heterogeneity, with at least half of all observed O-glycan abundance being the extended, doubly sialylated core 2 O-glycan structure. The low variability of this O-glycome was further tested in the molecular dynamic simulations. Previously, O-glycosylation on CD52-Fc was putatively localised to either T8 or its neighbour, S9 (3). Specific O-glycosylation sites have previously been identified as critical to protein function and activity in other well studied glycoproteins (15, 30–32), and the molecular dynamic modelling informed by experimental data infers this for CD52 binding to HMGB1 Box-B. High resolution mass spectrometry with paired EtHcD fragmentation techniques was able to localise the site of O-glycosylation of soluble CD52-Fc (produced in Expi293 host cells) to T8. Informed dynamic modelling of the soluble CD52 interaction with HMGB1-Box B showed that a lack of sialylation of the O-glycan on T8 would prevent binding of the CD52 peptide to HMGB1 Box-B since contacts with the protein directly involve the anionic terminal sialic acids binding to the cationic amino acids. From this model, the measured low site O-glycan site occupancy of an average of 7.5 % O-glycosylated rCD52 then appears to be the limiting factor in the low level of immunosuppressive activity in Expi293 cells. HMGB1-CD52 binding has been reported to be essential for CD52 activity (2) and our modelling of the doubly sialylated extended core 2, α-2,3 sialylated O-glycan on T8 of the soluble CD52 peptide confirmed that these sialic acids are essential for peptide binding to HMGB1 Box B, specifically interacting with positively charged amino acids unique to HMGB1 Box B, R91, K140, K146, R157 (PDB identification). Of note, we have shown (2) that CD52-Fc does not bind to HMGB1 Box-A, which is seen by the modelling to assume the same structure as the HMGB1 Box-B binding cleft, but without having three positively charged amino acids available to interact with the anionic sialic acids in the binding cleft.

Terminal sialic acids of the N-glycans on CD52 were orientated towards the solvent front with the N-glycan on HMGB1 wrapping around the N3 CD52 glycan in the MD simulations. This further implies that the N-glycan of CD52 plays a different role to the O-glycan, with the primary role of the N-glycan being to interact with the downstream immunoregulator SIGLEC-10. While additive force fields like those used here can enhance glycan-glycan interactions (23) overestimating glycan-glycan contacts, these predictions appear to be in accord with what is currently experimentally known of the soluble CD52 interaction.

Here we propose that the HMGB1-CD52 complex formed by the interaction of the doubly α-2,3 sialylated core 2 O-glycan on T8 of the CD52 peptide with the HMGB1 Box B binding cleft, forces the highly sialylated CD52 N-glycans at site N3 of CD52 to extend outwards, allowing them to interact with SIGLEC10, thus initiating immunosuppressive activity.

In conclusion, we were able to characterise both the N-glycome and O-glycome of recombinant soluble CD52 using glycopeptide and released glycomics mass spectrometry and have identified the probable most active glycoforms of soluble CD52. The O-glycosylation site of CD52 has been confidently localised to T8 for the first time. This deep characterisation of CD52 extends the current knowledge we have on the glycosylation of this therapeutically relevant glycopeptide. Experimental data obtained in this study then informed a molecular dynamics simulation which was able to infer a bi-modal mechanism of action involving both the O-glycans and N-glycans of CD52. The N- and O-glycans are shown to contribute to different interactions, with extended core 2 sialylated O-glycosylation needed to bind to HMGB1-Box B, while heavily sialylated N-glycosylation is needed to activate SIGLEC receptors (22), that mediate immune suppression. Our findings further define the structural basis for the bioactivity of CD52 and will assist in the glycoengineered production of CD52 as a therapeutic agent.

## Methods

### Recombinant CD52-Fc production

CD52-Fc was expressed as previously described (5). DNA encoding signal peptide and CD52 sequences were joined to human IgG1 Fc via PCR, then ligated into pCAGGS vector for transfection into Expi293 cells. The construct included a flexible GGSGG linker, a cleavage site for Factor Xa protease between the signal peptide and Fc molecule, a C-terminus strep-tag II sequence for purification (31; Supporting S10). CD52-Fc was purified from growth medium by affinity chromatography on Streptactin resin and eluted with 2.5mM desthiobiotin (5).

### MonoQ anion exchange column fractionation and protein quantitation

After recombinant CD52 production, samples were separated as previously described (3). In short, rCD52-Fc was diluted into 5 mL 50mM Tris-HCl, pH 8.3, and applied to a Mono Q column (Mono Q 5/50 GL, 10 µm particle size, 5 x 50 mm column dimensions, 1 mL column volume, 2 mL/min flow rate (GE Lifesciences, Parramatta, Australia). The column was washed with 10 column volumes of 50mM Tris-HCl, pH 8.3, and then eluted with 50 column volumes of 50mM Tris-HCl, 500mM NaCl, pH 8.3 in 0.5mL fractions. Protein content of eluted fractions was estimated using a Qubit protein assay (ThermoFisher Cat: Q33212).

### ELISpot assay of CD52-Fc activity

CD52-Fc MonoQ fractions were tested by interferon gamma (IFN-γ) ELISpot assay to measure immune suppression in peripheral blood mononuclear cells (PBMCs), as previously described (3). After approval by Melbourne Health Human Ethics Committee, blood samples were obtained with informed consent from healthy males. PBMCs were isolated from fresh blood on Ficoll/Hypaque (Amersham Pharmacia, Uppsala, Sweden), washed in phosphate-buffered saline (PBS) and re-suspended in Iscove’s Modified Dulbecco’s medium (IMDM) containing 5% (v/v) pooled, heat-inactivated human serum (PHS; Australian Red Cross, Melbourne, Australia), 100mM non-essential amino acids, 2 mM glutamine, and 50 μM 2- mercaptoethanol (IP5 medium). PBMCs (10^4^/well) were incubated in 200 μl IP5 medium in replicates of three in 96-well ELISpot plates (MultiScreen HTS, Millipore, Bayswater, VIC, Australia) for 18-24 h at 37° C in 5% CO2 air. Wells had been conditioned by washing with 35% ethanol before being coated with anti-human IFN-γ mAb (10 μg/ml) in PBS overnight at 4° C. PBMCs were incubated with anti-CD3/28 Dynabeads (1 bead/cell) +/- CD52-Fc fractions (5 ng/ml). After 24 h, cells were lysed with water and discarded. Wells were washed with PBS between sequential incubations with biotinylated anti-human IFN-γ (1ug/ml), streptavidin-alkaline phosphatase (Mabtech) and 5-bromo-4-chloro-3- indolyl-phosphate/nitro blue tetrazolium substrate solution (Mabtech). The colour reaction was stopped by addition of water and IFN-γ spots counted with an AID ELISpot Reader (Autoimmun Diagnostika Gmbh, Strassberg, Germany).

### Removal of Fc fragment from CD52-Fc

CD52 - Fc fusion protein (20µg) was cleaved with 2 µL of FXa protease (New England Biolabs, Ipswich, USA) in a volume of 350 µL of water containing 2 mM of CaCl2. Following cleavage, samples were applied to a C18 spin tip made from a frit of Empore C18 SPE disc (cat #66883U) and CD52 glycopeptide without Fc was eluted with 50 µL of 85% (v/v) acetonitrile (ACN) and the sample dried in a vacuum concentrator.

### Bottom-up MS/MS analysis of purified rCD52 peptide

Dried, purified Fxa-cleaved CD52 peptide was resuspended to an approximate concentration of 3 µg/µL equivalent of initial measured concentration and analysed on an Orbitrap Eclipse Tribrid Mass Spectrometer (Thermo Fisher Scientific) coupled to a Dionex Ultimate 3000 RSLC nano System (Thermo Fisher Scientific). Peptides were separated on an in-house packed C18 column (75 µm x 15 cm, 2 µm particle size, ReproSil-Pur 120 C18-AQ, Dr. Maisch). For this purpose, a gradient of (0-2 min—0 %, 3 min – 2 %, 5 min – 20 %, 25 min – 30 %, 30 - 40 min - 98 %, 43 - 50 min – 2 %) solvent B (99.9% ACN containing 0.1% (both v/v) formic acid (FA) with solvent A consisting of 0.1% (v/v) aqueous FA was used at a constant flow rate of 300 nL/min. Full MS scans were acquired in the range *m/z* 350-2,000 using 120,000 resolution, Automatic Gain Control (AGC) of 5 x 10^5^ ions and 100 ms maximum injection time. The mass spectrometer was operated in positive ion polarity mode. Data-dependant acquisition (DDA) was used to collect electron transfer higher-energy collision dissociation (EThcD)-MS/MS data within a 3 s cycle time. EThcD-MS/MS of precursors isolated with a quadrupole isolation window of *m/z* 2 was performed with a supplemental activation energy of normalised collision energy (NCE) 25%. Fragment ions were detected in the Orbitrap using 60,000 resolution, AGC of 2 x 10^5^ ions and a 120 ms maximum injection time.

### PNGase F release and O-glycan release

N-glycans were released from MonoQ fractionated FXa-cleaved CD52 using PNGase F (Promega). Briefly 10 U of PNGase F in MilliQ water was added to dried samples, then incubated at 37 degrees overnight. Following PNGase F release, samples were added to a self-packed C18 spin column made from a frit of Empore C18 SPE disc (cat #66883U) and pre-equilibrated with 0.1% FA. The flowthrough containing N-glycans was collected for reduction and analysis of the eluted N-glycans after carbon SPE cleaning, described below. The de-N glycosylated peptides retained on the C18 spin column (de- N-CD52 peptides) were eluted with 50 µL of 85% ACN, dried, then resuspended in deionized water to an equivalent concentration of 1 µg/µL of rCD52. O-glycans were released from de-N-CD52 peptides using β elimination with 0.5 M NaBH4 in 50 mM KOH for 16 hours. Following release, O-glycans were cleaned as described below, then resuspended for negative mode PGC (porous graphitised carbon) LC-MS/MS analysis, also described below.

### PGC cleaning of released N- and O-glycan alditols

Released N- and O-glycans were cleaned as previously described with modifications (33). Briefly, N-glycans were resuspended in 20 µL of 1 M NaBH4 in 50 mM KOH and incubated at 50 °C for 3 hours. After this incubation, all samples were diluted to 120 µL total volume with deionized water, then acidified with 2 µL of glacial acetic acid. Following acidification, N-glycan and O-glycan samples were both taken for cleaning of carbon solid phase extraction (SPE) as described (33). After cleaning, samples were dried, then resuspended in 9 µL of MilliQ for negative mode PGC LC-MS/MS analysis.

### Negative mode PGC-LC-ESI-MS/MS of released glycan alditols

8 µL of purified released N-and O- glycans was injected onto a Thermo HyperCarb PGC column (3 μm particle, 1 mm X 30 mm) using an Agilent 1260 HPLC coupled with a Thermo LTQ Velos Pro linear ion trap mass spectrometer. Glycans were chromatographically resolved over 60 min at a flow rate of 15 μL/min at 50 °C with the following gradient: Buffer A: 10 mM ammonium bicarbonate, Buffer B: 70 % (*v/v*) acetonitrile in 10 mM ammonium bicarbonate. The gradient parameters were: 0-3 min—0 % B, 4 min – 14 % B, 40 min – 40 % B, 48 min – 56 % B, 50 - 54 min – 100 % B, 56 – 60 min – 0 % B. The mass spectrometer was operated in negative ion mode and configured to perform one full zoom scan MS experiment (HESI source temperature 55 °C, spray voltage 2.75 kV, sheath gas flow 13, auxiliary gas flow 7, capillary temperature 275 °C, scan range 500-2000 *m/z*, AGC of 3e4, 3 microscans, and maximum IT of 100 msec) with the top 5 precursors (dynamic exclusion window of 15 sec) selected for MS/MS (scan range 150-2000 *m/z*, AGC of 1e^4^, maximum IT of 100 msec, isolation window 1.4 *m/z*, and normalized collision energy set as 33).

### Data analysis and search settings

Released glycan analysis was performed manually using the following criteria: *m/z* signals corresponding to biosynthetically possible glycan compositions according to GlycoMod (34) were selected for area-under-curve (AUC) quantitation using Skyline (35). Each peak area was then expressed as a percentage of the total area of all glycans in the sample. Structural characterisation was performed for glycans with MS/MS fragmentation using diagnostic ions (13, 36) as well as known PGC elution patterns of specific glycan features (37, 38).

Glycopeptide samples were searched using both MSFragger (39) (v20.0) with the following search settings: a custom FASTA protein database was used including common decoy contaminants provided from MSFragger and CD52 proteins identified in Supporting Information 4; glycan search list was defined by observed compositions from N-glycan and O-glycan analysis (Supporting Information S1, S3) and a cutoff Pep2D filter of < 0.001 was applied. MS2 search parameters were set according to Thermo Scan headers, and all search tolerances were set to 20 ppm. MSFragger searching used the default Glyco-O-Hybrid workflow with a combined composition list of N-glycans and O-glycans analytically identified (Supporting Table S1, S3). Site occupancy of N- and O- glycosites was calculated using a targeted mass list in Skyline by comparing the AUC of non-glycosylated CD52 fragments to deamidated CD52 fragments (Supporting Information S6).

### Modelling of interactions between CD52 and HMGB1-Box B

To understand the molecular determinants for recognition of the CD52 glycopeptide by HMGB1 Box B, and the role of sialic acid in this interaction, a series of all-atoms, classical MD simulations based on deterministic sampling was run, first on the isolated glycopeptide, then on the CD52/HMGB1 Box B complex which was built based on information gathered from the simulations of the isolated CD52 glycopeptide. To build the starting structure of the CD52 peptide the aa 25-36 region was selected from the Uniprot P31358 CAMPATH-1 antigen (12 residue peptide with sequence GQNDTSQTSSPS) from the AlphaFold Protein Structure Database (https://alphafold.ebi.ac.uk/). This sequence is recognised and bound by the HMGB1 Box B and corresponds to the functional part of the peptide used in the experimental assays in this work. The glycan structures were chosen based on the results of the released glycome and glycoproteome analysis in this work. To represent the fully sialylated CD52 glycopeptide, a core-fucosylated tetra-antennary α-2,3 tetra-sialylated N-glycan (GlyTouCan ID G80552MJ) was linked at position N3, and a α-2,3 di-sialylated core 2 oligosaccharide (GlyTouCan ID G42089IU) at position T8. For the reconstruction of the peptide’s glycosylation, the glycan 3D structures were sourced from the GlycoShape Glycan Database (https://glycoshape.org) and used the ReGlyco tool (21) to link the N- and O-glycans at positions N3 and T8, respectively. Two sets of MD simulations of the isolated CD52 were run, one with the peptide linked to fully sialylated N- and O-glycans and the other where neither oligosaccharide had terminal sialic acids (Figure 4a and b). Five uncorrelated MD simulations of 1 μs production were run for each set, with a cumulative sampling of 10 μs for the CD52 glycopeptide in the presence and absence of sialic acid. The starting structures described above were used for the first MD simulation in each set, while the remaining trajectories were started from uncorrelated snapshots taken from these MD trajectories.

The system set-up consisted of an initial energy minimization phase of 500,000 steps of steepest descent. The system was then brought at room temperature through a two-step heating scheme in the NVT ensemble of 500 ps each. An integration time step of 2 fs was used throughout. The temperature was raised from 0 K to 100 K during the first step and from 100 K to 300 K during the second step through Langevin dynamics with a friction coefficient (gamma_ln) set to 1.0 ps−1. The heating phase was followed by a 500 ps NPT equilibration to bring the system into the isothermal-isobaric (NPT) ensemble at 1 atm. Pressure was controlled with a Berendsen barostat in the simulations run with version 2018 of the AMBER package (40) and with the Parrinello-Rahman in the simulations run with version 2021.4-Ubuntu-2021.4-2 of GROMACS (41). Long range electrostatics were treated with Particle Mesh Ewald (PME) with an 11 Å cutoff. Long range dispersion interactions were truncated with an 11 Å cutoff. In all simulations, the GLYCAM_06j-1 force field (42) was used to represent the carbohydrate atoms, the TIP3P model (43) was used to represent the solvent, and ff14SB parameters were used to represent the peptide and counterions (44). Simulations were run with both AMBER 2018 on GPU (45) and with 2021.4-Ubuntu-2021.4-2 GROMACS (46) on ORACLE Cloud Infrastructure (OCI).

The complex between the fully sialylated CD52 glycopeptide and HMGB1 was obtained by structural alignment of the dominant conformation at equilibrium of the CD52 backbone, obtained from Ramachandran analysis of the backbone, to the 12 aa sequence SQAMDDLMLSPD of the p53 transactivation domain in complex with HMGB1 Box A (PDB-ID 2LY4) with pymol (www.pymol.org).

This p53 transaction domain complex had been previously modelled and was used for starting structural alignment. After structural alignment of the peptides, the starting structure for the complex was obtained by replacing the structure of HMGB1 Box A with the structure of Box B from PDB 2GZK, from which we removed the previously modelled bound DNA and the Box A domain. The MD simulation of this complex was set-up following the protocol described above and the production trajectory was run for 1 μs.

## Data Availability

Raw files relating to released glycans for this work are accessible on GlycoPost (Accession ID GPST000491 with reviewer pin 1669) (47), with glycoproteomic data accessible via ProteomeXchange (PXD056562)(48).

## Supporting information

This article contains supporting information.

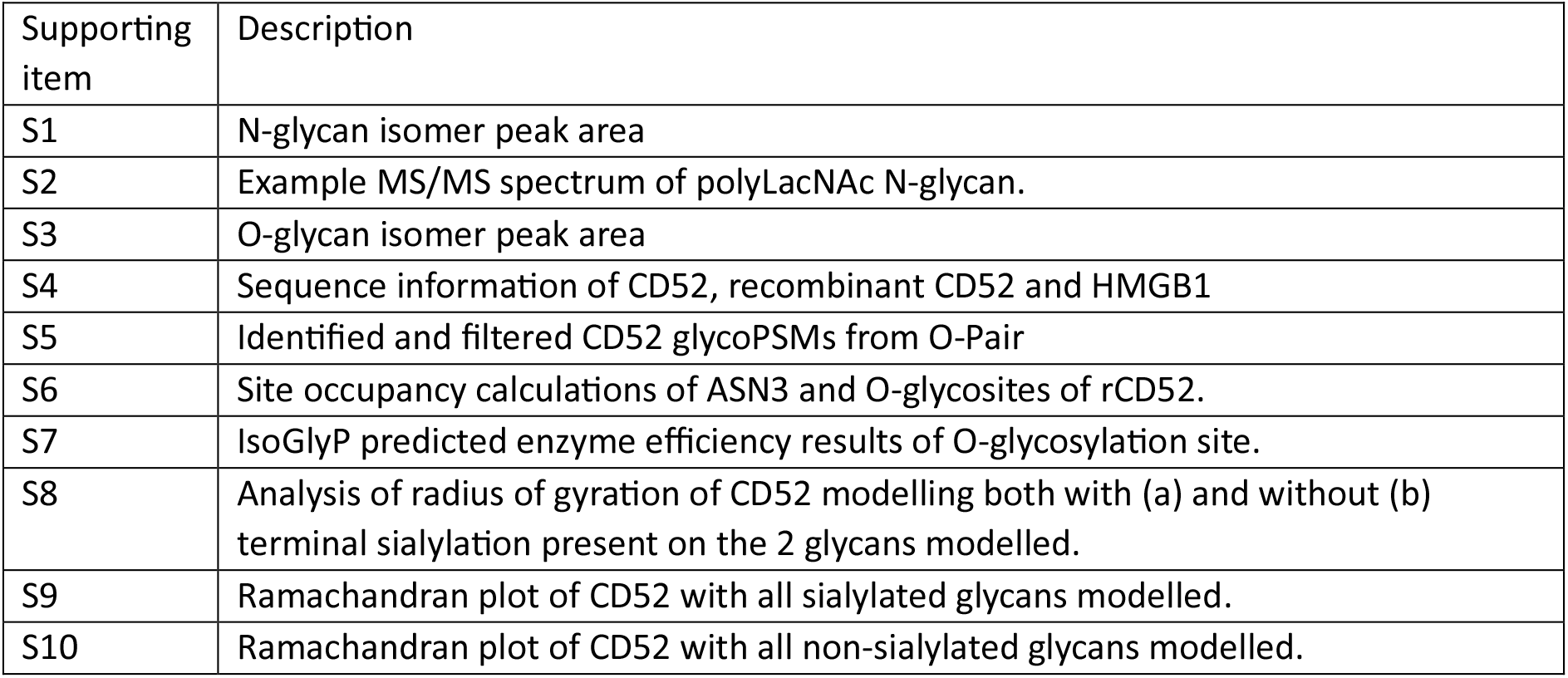

Supporting items S1, S3, S4, S5, S6, and S10 are in attached .xlsx file.

## Author Contributions

N.D., S.D., E.B.S, M.Z, E.S.X.M, E. G-B and E.F. all contributed to experimental work. N.D., S.D., E.B.S, M.Z, E.S.X.M, E.F., L.C.H. and N.P. all assisted in data analysis and interpretation. N.D., S.D., M.Z., E.S.X.M, E.F., L.C.H. and N.P. contributed to writing and editing the manuscript.

## Acknowledgements

N.D. was supported by a Macquarie University Research Excellence Scholarship (MQRES) and a CSIRO Future Science Platform Top-Up Scholarship. E.S.X.M. was supported by Australian Research Council Centre of Excellence in Synthetic Biology (CE200100029). LCH was supported by a NHMRC Leadership Investigator Grant (APP1173945), a NHMRC Program Grant (APP1150425), Victorian State Government Operational Infrastructure Support and the NHMRC Research Institute Infrastructure Support Scheme. Some of the research described herein was facilitated by access to the Australian Proteome Analysis Facility (APAF) funded under the Australian Government’s National Collaborative Research Infrastructure Strategy (NCRIS)/Education Investment Fund.

