## Supporting Information S2, 8, 9, 10 for "The molecular basis of immunosuppression by soluble CD52 is defined by interactions of N-linked and O-linked glycans with HMGB1 Box B"

**Soluble CD52 activity is modulated by its N-linked and O-linked glycans**

^1^ARC Centre of Excellence in Synthetic Biology, School of Natural Sciences, Macquarie University, Sydney, Australia, ^2^Department of Chemistry, Maynooth University, Maynooth, Ireland, ^3^The Walter and Eliza Hall Institute of Medical Research, Melbourne, VIC, Australia ^4^Australian Proteome Analysis Facility, Macquarie University, Sydney, NSW, Australia; ^5^School of Biological Sciences, University of Southampton, Southampton, United Kingdom

^+^Current Address: Institute for Biomedicine and Glycomics, Griffith University, Southport, QLD, Australia

**KEY WORDS:** CD52, HMGB1, glycomics, glycoprotein, mass spectrometry, molecular dynamics

**Subject category:** Analytical Glycobiology

**Supporting information**:

| Supporting item | Description |
| --- | --- |
| S1 | N-glycan isomer peak area |
| S2 | Example MS/MS spectrum of polyLacNAc N-glycan. |
| S3 | O-glycan isomer peak area |
| S4 | Sequence information of CD52, recombinant CD52 and HMGB1 |
| S5 | Identified and filtered CD52 glycoPSMs from O-Pair |
| S6 | Site occupancy calculations of ASN3 and O-glycosites of rCD52. |
| S7 | IsoGlyP predicted enzyme efficiency results of O-glycosylation site. |
| S8 | Analysis of radius of gyration of CD52 modelling both with (a) and without (b) terminal sialylation present on the 2 glycans modelled. |
| S9 | Ramachandran plot of CD52 with all sialylated glycans modelled. |
| S10 | Ramachandran plot of CD52 with all non-sialylated glycans modelled. |

Supporting items S1, S3, S4, S5, S6, and S10 are in attached .xlsx file.

**S2	Example MS/MS spectrum of tri-antennary, tetra-sialylated N-glycan containing a di-sialic acid.**

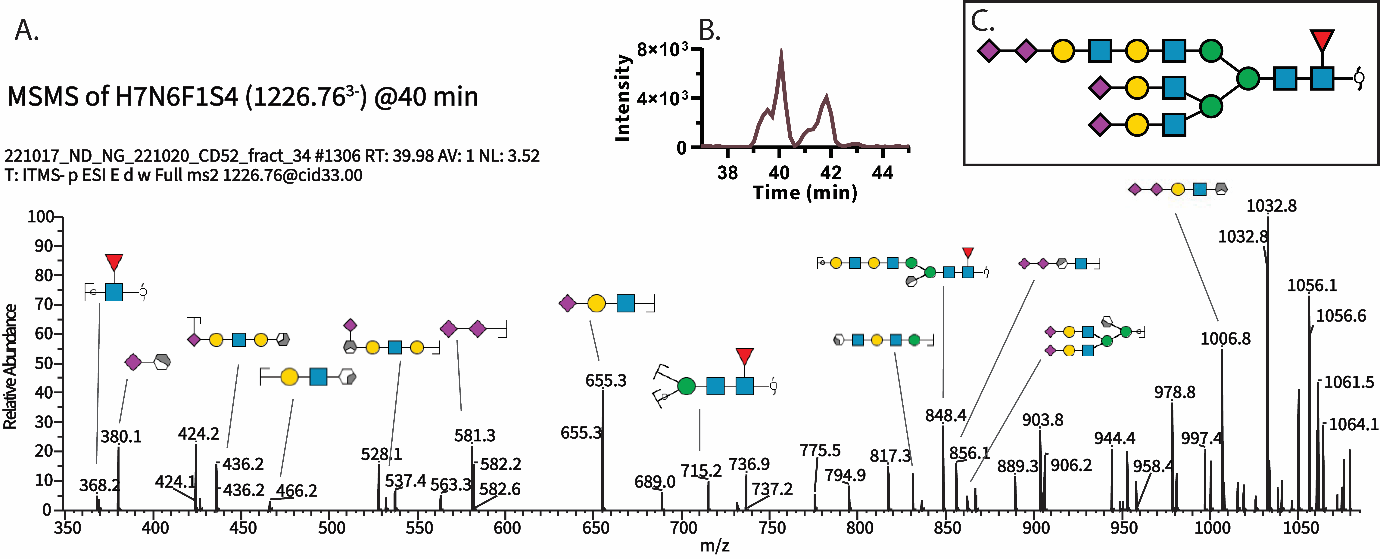

A) MS/MS analysis of N-glycan isomer H7N6F1S4 (1226.76^3-^). Several clear diagnostic glycan signatures provide clues as to the structure of this glycan, including m/z 368.2 (core fucosylation), 436.2 (sial-LacDiNAc extension), and 581.3 (di-sialylation) among others. B) The elution profile of this glycan composition shows separation of two major isomers. It is possible that there are other isomers in these peaks which have not been sufficiently chromatographically resolved. C) Solved structure of the H7N6F1S4 glycan isomer eluting as the 40-minute peak.

**S8	Analysis of radius of gyration of CD52 modelling both with (a) and without (b) terminal sialylation present on the 2 glycans modelled.**

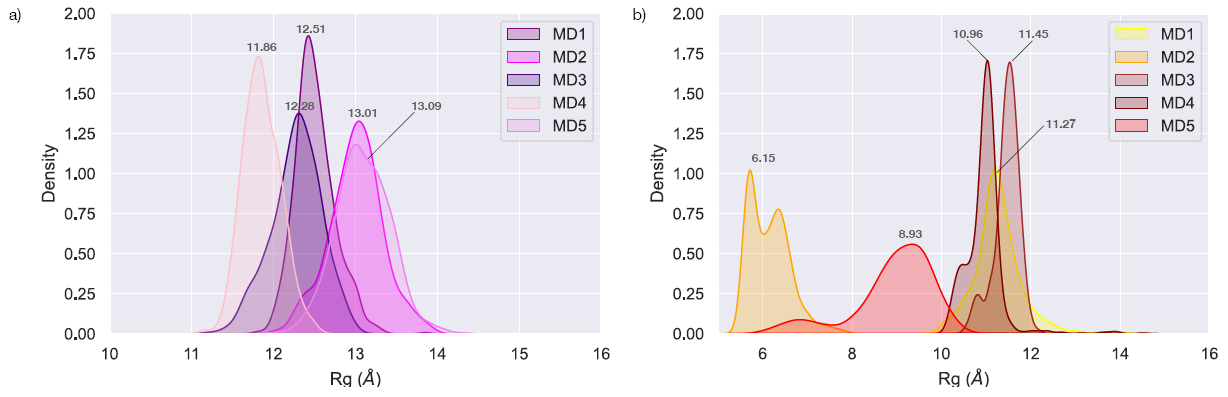

Analysis of radius of gyration of CD52 modelling both with (a) and without (b) terminal sialylation present on the 2 glycans modelled. Modelled glycans included an N-glycan on N3 (GlyTouCan ID G80552MJ or GlyTouCan ID G56655CC) and an O-glycan on T8 (GlyTouCan ID G96017QA or GlyTouCan ID G42089IU). With terminal siaylation present, glycosylated CD52 was modelled to be have a more uniform, extended position than CD52 that was lacking terminal sialylation.

**S9	Ramachandran plot of CD52 with all sialylated glycans modelled.**

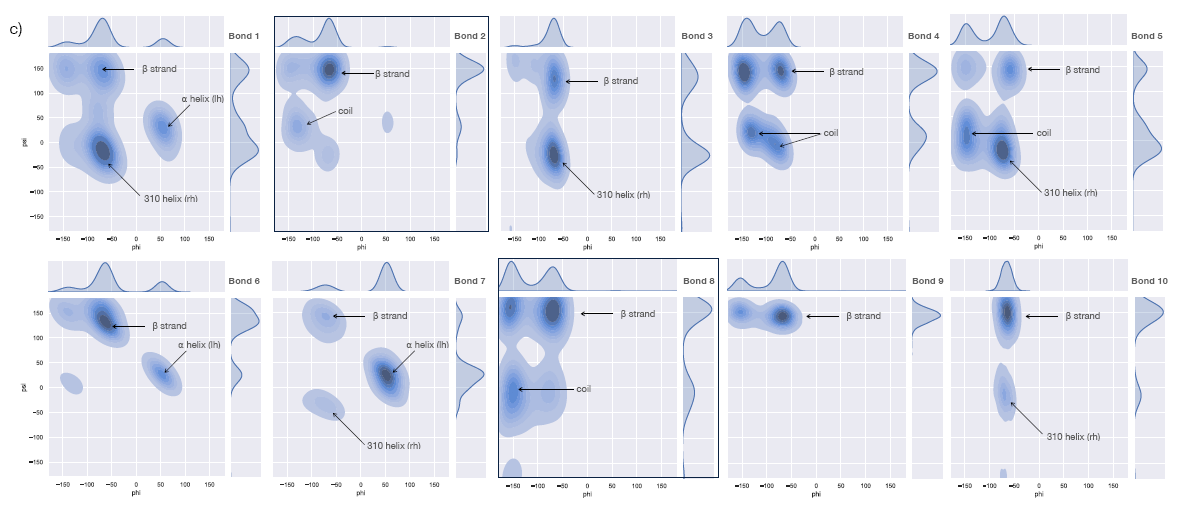

Ramachandran plot of CD52 with all sialylated glycans modelled (GlyTouCan ID G42089IU on T8 and GlyTouCan ID G80552MJ on N3).

**S10	Ramachandran plot of CD52 with all non-sialylated glycans modelled.**

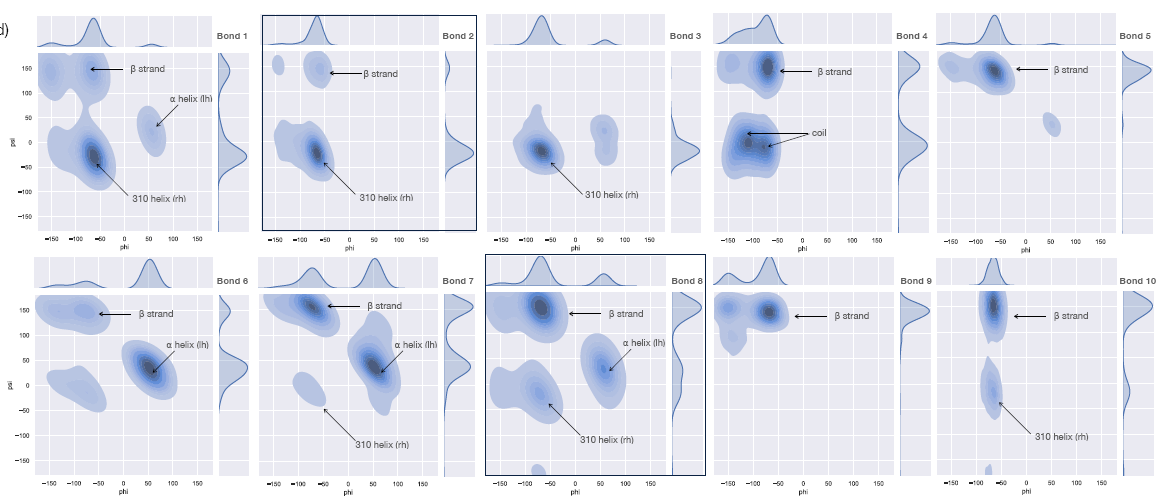

Ramachandran plot of CD52 with all non-sialylated glycans modelled (GlyTouCan ID G96017QA on T8 and GlyTouCan ID G56655CC on N3).
